# Virtual Growing Child (VGC): A general normative comparative system via quantitative dynamic MRI for quantifying pediatric regional respiratory anomalies with application in thoracic insufficiency syndrome (TIS)

**DOI:** 10.1101/2024.04.28.591554

**Authors:** Yubing Tong, Jayaram K. Udupa, Joseph M. McDonough, Lipeng Xie, You Hao, Yusuf Akhtar, Caiyun Wu, Chamith S. Rajapakse, Samantha Gogel, Oscar H. Mayer, Jason B. Anari, Drew A. Torigian, Patrick J. Cahill

## Abstract

**Background:** A normative database of regional respiratory structure and function in healthy children does not exist.

**Methods:** VGC provides a database with four categories of regional respiratory measurement parameters including morphological, architectural, dynamic, and developmental. The database has 3,820 3D segmentations (around 100,000 2D slices with segmentations). Age and gender group analysis and comparisons for healthy children were performed using those parameters via two-sided t-testing to compare mean measurements, for left and right sides at end-inspiration (EI) and end-expiration (EE), for different age and gender specific groups. We also apply VGC measurements for comparison with TIS patients via an extrapolation approach to estimate the association between measurement and age via a linear model and to predict measurements for TIS patients. Furthermore, we check the Mahalanobis distance between TIS patients and healthy children of corresponding age.

**Findings:** The difference between male and female groups (10-12 years) behave differently from that in other age groups which is consistent with physiology/natural growth behavior related to adolescence with higher right lung and right diaphragm tidal volumes for females(p<0.05). The comparison of TIS patients before and after surgery show that the right and left components are not symmetrical, and the left side diaphragm height and tidal volume has been significantly improved after surgery (p <0.05). The left lung volume at EE, and left diaphragm height at EI of TIS patients after surgery are closer to the normal children with a significant smaller Mahalanobis distance (MD) after surgery (p<0.05).

**Interpretation:** The VGC system can serve as a reference standard to quantify regional respiratory abnormalities on dMRI in young patients with various respiratory conditions and facilitate treatment planning and response assessment.

**Funding:** The grant R01HL150147 from the National Institutes of Health (PI Udupa).

## Introduction

It is important to collect quantitative measurements of regional respiratory structure and function from healthy children to serve as a reference standard for comparison with pediatric patients with respiratory disease for disease characterization, treatment planning, and response assessment. This is because there are many scenarios in clinical practice where utilization of corresponding normative measurements obtained in healthy children has been shown to be useful. For example, trunk motion comparisons between children with moderate adolescent idiopathic scoliosis (AIS) and healthy children in [1]; bone age comparisons between patients with bone disorders and healthy children were illustrated in [2]; cognitive function comparisons between children with brain tumors and healthy children in [3]; urinary phosphorus excretion comparisons between children with urolithiasis and healthy children in [4]; vitamin D from plasma comparisons between children with celiac disease and healthy children in [5]; and liver and spleen volume from CTs were collected from healthy children for providing reference ranges and potential thresholds to identify liver and spleen size abnormalities that might reflect disease in children [6].

Yet, such a large open-source normative dataset in relation to regional respiratory structure and function does not exist. Without such as normative database, it is not possible to quantify the distribution and severity of abnormalities that may exist in a patient with respiratory dysfunction. For example, in pediatric patients with thoracic insufficiency syndrome (TIS), a serious childhood disorder related to the inability of the thorax to support normal respiration or lung growth [24,26], it is important to determine the locations and amounts of abnormality in respiratory structure and function before surgery so that optimal treatment strategies may be implemented, as well as to ascertain the actual effects of therapeutic intervention once performed.

One early study used longitudinal data sets to track lung function in healthy children and adolescents in order to understand better the growth of normal lung function over different phases of life [7]. By checking the normal lung index on CT (a ratio of lung volume in a range of densities considered to be normal (−950 to −700 HU) related to the total lung volume (−1024 HU to −250 HU) at full inspiration), clinicians can make an estimation of obstructive and restrictive lung disease [8, 9]. Pulmonary function testing (PFT) was proposed decades ago [10,11] and is still widely used as the reference standard for evaluating lung function [7-9,12-16]. However, PFT is useful only for evaluating subjects who are cooperative and can follow instructions during testing [17,18]. Moreover, PFT cannot determine the regional ventilation or the portion of exhaled lung volume from diaphragmatic and chest wall component displacements, let alone from the left lung separately from the right lung [19-21].

Although some other methods to check maximum respiratory pressures [22] as well as electrical impedance tomography [23] might be useful to detect changes in lung parenchymal aeration, they do not provide any anatomical information. Considering the temporal and spatial resolutions necessary for assessing lung anatomy, function, and dynamics, quantitative dynamic magnetic resonance imaging (QdMRI) provides a practical solution [24-26]. QdMRI allows for free-breathing image acquisitions and can measure respiratory dynamics in terms of volumes and tidal volumes of the lungs, hemi-diaphragms, and hemi-chest walls [26]. The accuracy of QdMRI in terms of volume measurement has been reported as ∼97% using a 4D dynamic phantom and one adult with breathing control during image acquisition [24].

Other imaging methods for evaluating lung structure or function like CT [31] which has the radiation concerns, single plane 2D dynamic MRI not covering 3D lungs [30], hyperpolarized gas MRI [27-29, 31-33] which may not be feasible for young patients in clinical practice. Some other methods need patient cooperation such as breath holding [34, 35]. Compared with ultrashort echo time (UTE) MRI [36] and phase-resolved functional lung (PREFUL) MRI [37], QdMRI provides a more practical solution with higher spatial resolution and comparable temporal resolutions. UTE MRI with limited field of view was used for checking lung function, only on adult study and much elder children than our TIS patients [38]. More importantly, QdMRI allows one to analyze the properties of multiple objects of interest beyond the lungs alone (e.g., left and right lungs, left and right hemi-diaphragms, liver, left and right kidneys, spleen, etc.) including those related to architecture and dynamics simultaneously, which is otherwise not feasible through other existing techniques.

The purpose of the current study is to introduce a system based on dMRI from the ongoing virtual growing child (VGC) project, which includes both healthy children data sets and corresponding quantitative normative reference standard measurements and models that can be used for analyzing respiratory anomalies and improving treatment planning. In particular, one aim of the VGC project is to use information gained from QdMRI, including 4D dynamics of the lungs, hemi-diaphragms, and hemi-chest walls, to facilitate surgical planning and response assessment in pediatric patients with TIS, which to our knowledge has not been previously reported. This approach can also be extended to other pediatric and adult conditions in the future.

The VGC database with dMRI images and corresponding object segmentations can also be useful for further artificial intelligence (AI) and machine learning based research on dMRI-based object segmentation and analysis. The database has 3,820 3D segmentations (around 100,000 2D slices with segmentations), which to our knowledge is unique and the largest dMRI dataset of healthy children, and which will be available publicly in the future.

## 1. Materials & Methods

### Image data sets and study subjects

This prospective study was conducted following approval from the Institutional Review Board (IRB) at the Children’s Hospital of Philadelphia, the single IRB of record, along with a Health Insurance Portability and Accountability Act waiver.

We enrolled 368 healthy children 6 to 14 years of age with the following exclusion criteria: (1) history of thoracic surgery, (2) history of asthma or other lung disease, (3) respiratory tract illness within the previous 30 days, and (4) history of scoliosis or other congenital skeletal abnormality. An image review process excluded low-quality MRI scans and images representing irregular breathing or body movement during acquisition.

In total, dMRI scans were acquired in 191 healthy children and then constructed with a 4D image per subject, and 3D volumes at end-inspiratory (EI) and end-expiratory (EE) time points were segmented with each having 10 object segmentations, leading to a total of 3,820 (191×2×10) 3D segmented object samples. Each object sample has a 3D segmentation mask covering an average of 25-30 slices, for a total of 95,500-114,600 2D slices with segmentations in the database. Table 1 shows the age and gender information of this healthy child cohort.

**Table 1.**
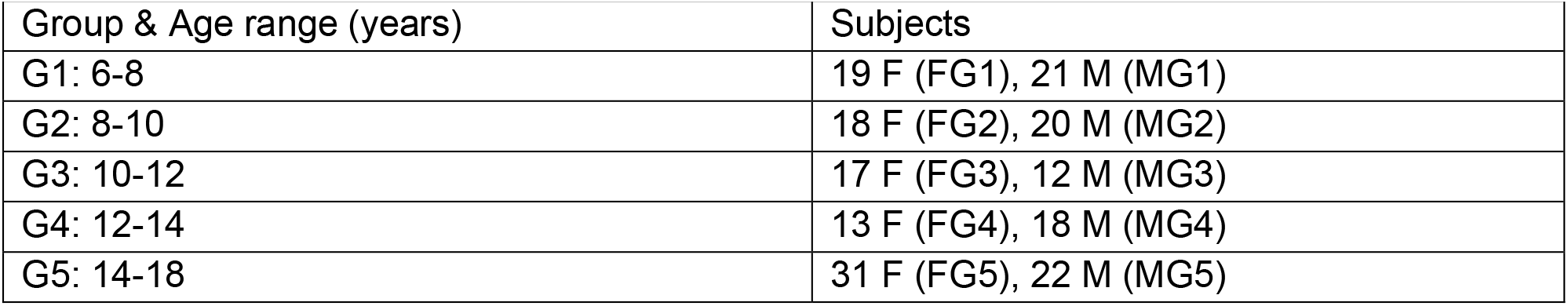
Demographic characteristics of VGC study healthy child cohort. VGC-virtual growing child; F-female, M-male, G-group, FG-female group, MG-male group.

|  |  |
| --- | --- |
| Group & Age range (years) | Subjects |
| G1: 6-8 | 19 F (FG1), 21 M (MG1) |
| G2: 8-10 | 18 F (FG2), 20 M (MG2) |
| G3: 10-12 | 17 F (FG3), 12 M (MG3) |
| G4: 12-14 | 13 F (FG4), 18 M (MG4) |
| G5: 14-18 | 31 F (FG5), 22 M (MG5) |

To illustrate an application of the VGC model, we also collected dMRI data sets in 49 pediatric TIS patients, where each TIS patient underwent dMRI obtained before and after surgical treatment. Table 2 shows the age and gender information of this TIS patient cohort.

**Table 2.** Demographic characteristics of TIS patient cohort (mean ± SD). TIS-thoracic insufficiency syndrome, SD-standard deviation, F-female, M-male.

| Table 2. Demographic characteristics of TIS patient cohort (mean ± SD). TIS-thoracic insufficiency syndrome, SD-standard deviation, F-female, M-male. | | |
| --- | --- | --- |
|  | Pre-operative | Post-operative |
| 22 F | 3.7±3.5 years | 6.5±3.6 years |
| 27 M | 3.4±3.5 years | 5.4±3.6 years |
| 22 F + 27 M | 3.5±3.5 years | 5.9±3.6 years |

### Methods

The VGC system provides image data sets, object segmentations, and quantitative measurements from QdMRI techniques. The steps involved in QdMRI generally include data acquisition, image construction, image processing, object segmentation, and object parameter measurements as described briefly below.

i. Free-breathing thoracoabdominal dMRI data acquisition: The dMRI scan protocol was as follows: 3T MRI scanner (Verio, Siemens, Erlangen, Germany), true-FISP bright-blood sequence, TR=3.82 ms, TE=1.91 ms, voxel size ∼1×1×6 mm3, 320×320 matrix, bandwidth 258 Hz, and flip angle 76°. With recent advances [25], for each sagittal location across the thorax and abdomen, we acquire 40 2D slices over several breathing cycles at ∼480 ms/slice. On average, 35 sagittal locations are imaged, yielding a total of ∼1400 2D MRI slices, with the resulting total scan time of 11-13 minutes for any subject [39].
ii. 4D image construction: For the acquired dMRI scans, we utilized an automated 4D image construction approach [25] to form one 4D image over one breathing cycle (consisting of typically 5-8 respiratory phases) from each acquired dMRI scan to represent the whole thoraco-abdominal body region dynamically. The algorithm selects roughly 175-280 slices (35 sagittal locations × 5-8 respiratory phases) from the 1400 acquired slices in an optimal manner using an optical flux method [50].
iii. Image processing: Intensity standardization (IS) is a well-established technique that allows for generation of MRI standardized signal intensity (sSI) values to attain tissue-specific numeric meaning [40]. Intensity standardization is performed on every time point/3D volume of one 4D image. Via IS, one can quantify the aeration properties of lung tissue at EI and EE [41].
iv. Object segmentation: For each subject, there are 10 objects segmented at both EI and EE time points in this database. They include the thoracoabdominal skin outer boundary, left and right lungs, liver, spleen, left and right kidneys, diaphragm, and left and right hemi-diaphragms. All of the healthy children in this study underwent larger field of view (LFOV) dMRI imaging, which included the full thorax and abdomen in the sagittal dMRI images. The superior-most aspect of the thoracoabdominal body region is defined as 15 mm superior to the lung apices (top red line in Figure 1), and the inferior-most aspect of the thoracoabdominal body region is defined as the inferior aspect of the kidneys (bottom red line in Figure 1). We used a pretrained U-Net based deep learning network to segment each lung in the intensity-standardized 4D image with a mean and standard deviation (SD) of Dice Coefficient (DC) of 0.97 ± 0.02 [42]. The liver and kidneys were segmented by using our previous anatomy-guided deep learning approach, with DC of 0.91±0.02 for liver and 0.87±0.05 for kidneys [43]. All auto-segmentation results were visually checked and manually refined as needed, under supervision of a board-certified radiologist (DAT) with 25 years of expertise in MRI and thoracoabdominal radiology. Figure 2 shows some examples of object segmentations at EI and EE time points. Example of 3D renditions of objects including lungs, liver, kidneys, spleen, and hemi-diagrams are shown in Figure 3.
v. Object parameter measurements: Four categories of parameters are included in the quantitative measurements:
  1. Morphological parameters: Height, surface area, and volume of lungs and diaphragms (left and right separately) at EI and EE time points were measured.
  2. Architectural parameters: The architectural analysis involves assessment of the magnitudes at EI and EE of the 3 sides and 2 angles of the four triangles connecting (i) the apex of the right hemi-diaphragm dome and the centroids of the left and right lungs, (ii) the apex of the left hemi-diaphragm dome and the centroids of the left and right lungs, (iii) the apex of the right hemi-diaphragm dome and the centroids of the liver and right kidney, and (iv) the apex of the left hemi-diaphragm dome and the centroids of the right and left kidneys. The objects in the architectural analysis are shown in Figure 3 with an example of a triangle for left hemi-diaphragm, left kidney, and right kidney shown for illustration.
  3. Dynamic parameters: Tidal volumes of lungs, hemi-diaphragms, and hemi-chest walls (left and right separately) and architectural parameter changes from EI to EE time points were measured. Diaphragm motion from EI to EE is also computed by uniformly selecting points on the surface region of the diaphragm [39].
  4. Developmental parameters: Change in dynamic parameters from one age group to the adjacent higher age group.

**Figure 1.**
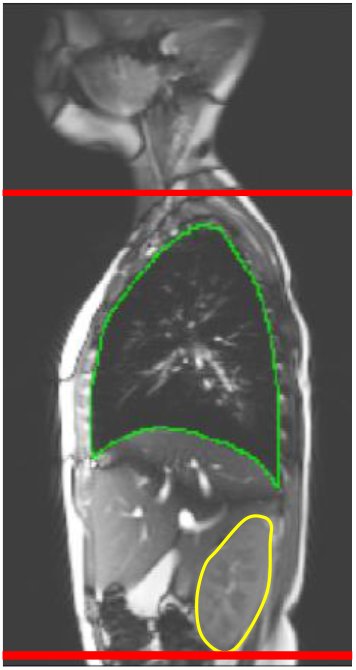
Thoracoabdominal body region illustration on sagittal dMRI.

**Figure 2.**
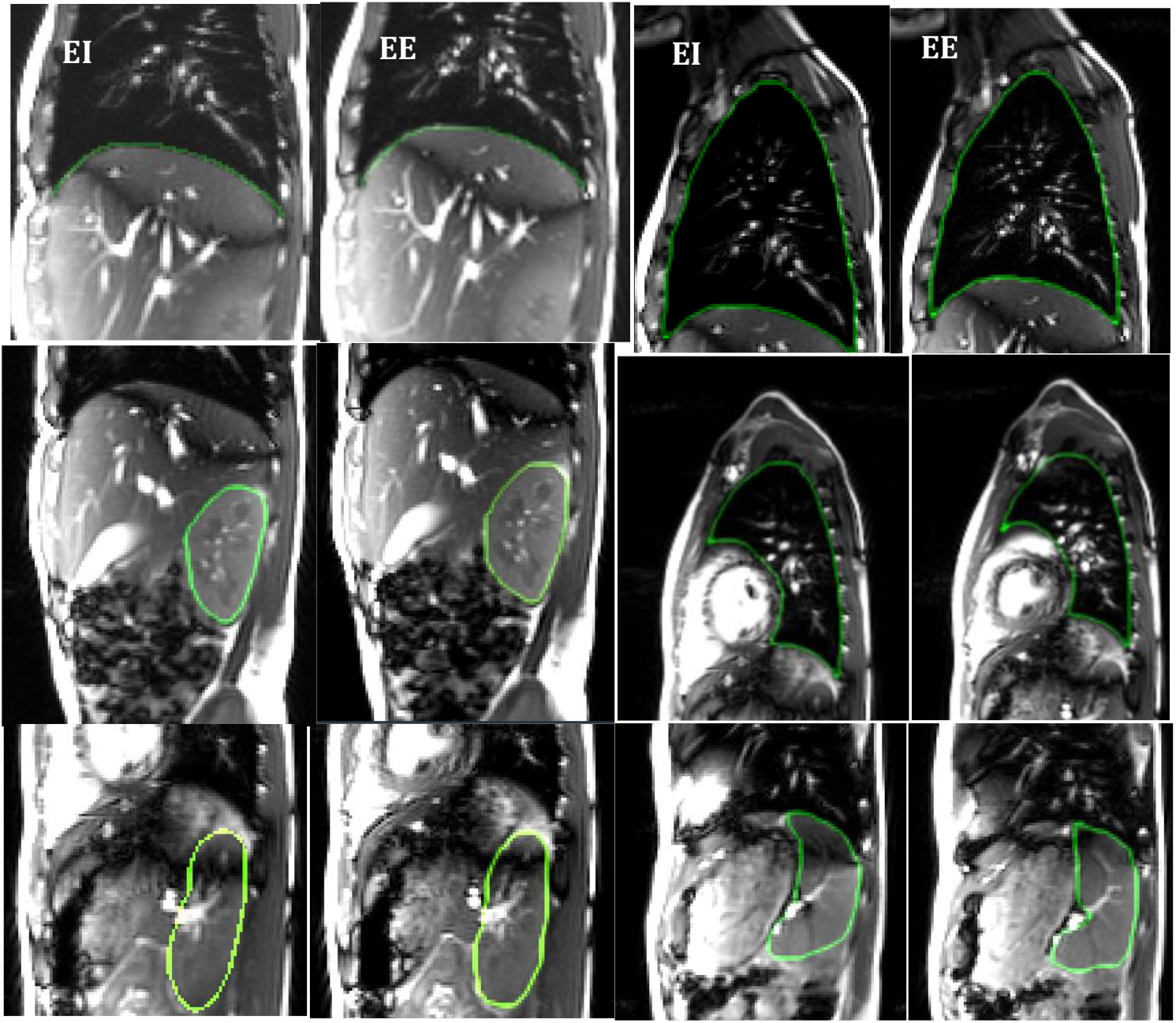
Examples of object segmentations at EI and EE including right hemi-diaphragm and right lung (top row); right kidney and left lung (middle row); left kidney and spleen (bottom row). EI: end inspiration, EE: end expiration.

**Figure 3.**
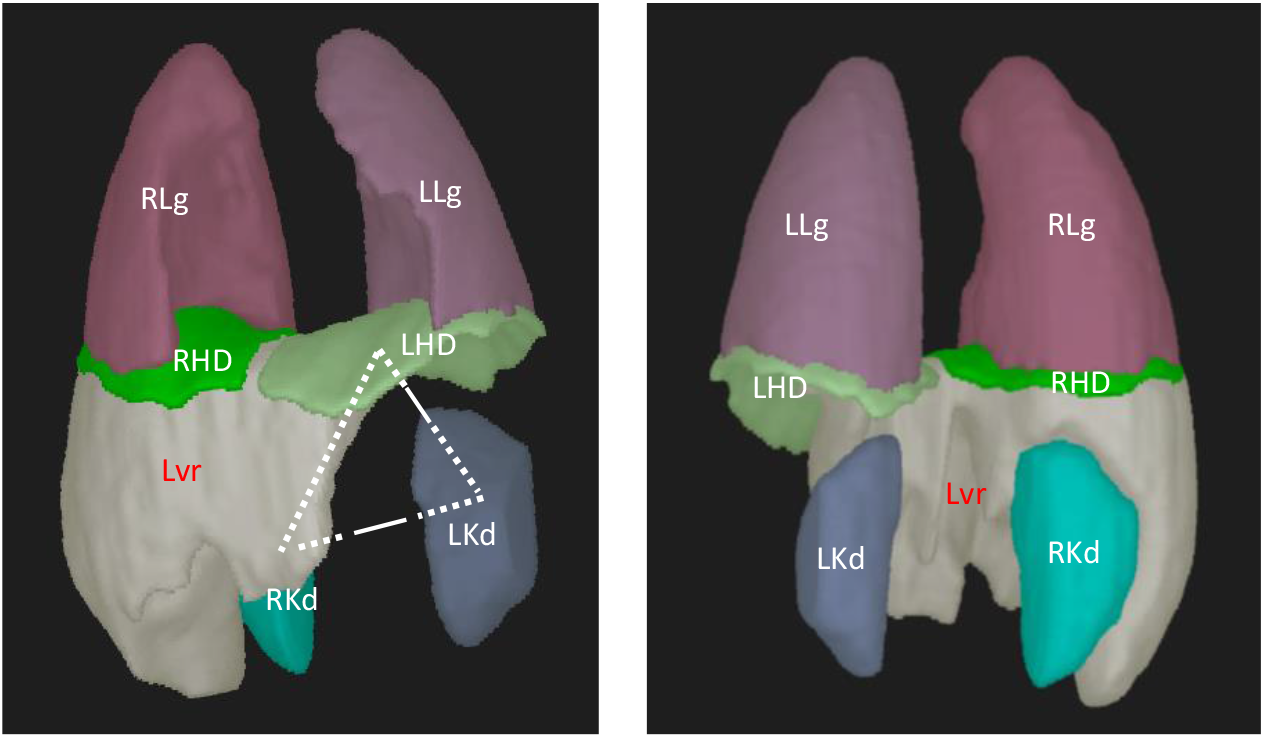
Objects in the architectural analysis shown in anterior to posterior view (left) and posterior to anterior view (right) for a normal healthy child. LLg/RLg = left/right lung, LHD/RHD = left/right hemi-diaphragm, Lvr = liver, LKd/RKd = left/right kidney.

In total, 103 quantitative parameter measurements are obtained in this study. The definitions of these measurements are listed in supplementary Table S1.

### Statistical analysis

The statistical toolbox in MATLAB (R2019b, MathWorks, Inc., Natick, MA) was utilized. Two kinds of analysis are performed based on the quantitative measurements as follows.

Age and gender group-based analyses and comparisons for the VGC study healthy child cohort for the quantitative parameter measurements described above. Two-sided unpaired t-testing was performed to compare mean measurements of left vs. right sides of bilateral objects, and mean measurements of objects at EI vs. EE for age and gender specific groups. We also assessed the Pearson correlation between QdMRI and pulmonary function testing (PFT) measurements in a subset of 51 healthy children from the VGC cohort who underwent both QdMRI and PFT. PFT is often used to measure global respiratory function including that of the lungs bilaterally. We therefore assessed the Pearson correlation between the bilateral lung volume at end-expiration (BLVee) derived from QdMRI and the functional residual capacity (FRC), the volume of air in the lungs following normal expiration, derived from PFT.

1) Application of VGC measurements/ models for assessment of the TIS patient cohort. First, an extrapolation approach is utilized to estimate the associations between object parameter measurements and age via a linear model. Then, the linear model is used to predict the measurements for very young TIS patients, given that the age of some of the patients is less than 6 years of age, which is the lower age limit of subjects in the VGC study healthy child cohort. Subsequently, the differences of object parameter measurements of individual TIS patients and those of corresponding age- and gender-matched healthy children are determined, in order to quantitatively estimate the deviations from normality in TIS patients before and after surgery to evaluate treatment effects. Mahalanobis distance (MD), a metric for measuring the statistical distance between a random sample x and the probability distribution of a random variable X, is used for this purpose. It is applicable even when X is an n-dimensional random vector. When n = 1, MD corresponds to the number of standard deviations of X that x is away from the mean of X (also known as Z-score). When X is a random vector with n elements and x is random sample vector with n elements, MD(x) has a similar meaning as in the scalar case, although now it considers the correlations among the elements of the random vector in estimating the distance.

## 2. Results

### QdMRI measurements and analysis on the VGC study healthy child cohort

Table 3 shows the global comparison between the male group (94) and female group (97) of healthy children in the VGC study. Five salient measurements among all 88 object parameter measurements, including left hemi-diaphragm height at EI (M1) and EE(M3), left hemi-diaphragm surface area at EI (M17), and the length between centroids of right and left kidneys at EI (M32) and EE (M41), show significantly larger values in the male group compared to the female group.

**Table 3.** Salient measurement comparisons between male (94) vs. female (97) healthy children.

QdMRI measurements and analysis on the VGC study healthy child cohort
|  |  |  |  |  |  |
| --- | --- | --- | --- | --- | --- |
| **Features** | **M1** | **M3** | **M17** | **M32** | **M41** |
| **Male (94)** | 4.69<br>0.98 | 4.99<br>1.09 | 293.48<br>74.71 | 10.49<br>1.54 | 10.42<br>1.44 |
| **Female (97)** | 42.89<br>8.77 | 4.54<br>1.05 | 272.78<br>61.08 | 10.06<br>1.13 | 9.89<br>1.08 |
| **P values** | <0.0001 | <0.0001 | 0.04 | 0.03 | <0.0001 |
| M1 - left hemi-diaphragm height at EI (cm); M3- left hemi-diaphragm height at EE (cm); M17 - left hemi-diaphragm surface area at EI (cm<sup>2</sup>); M32/M41 - distance between centroids of right kidney to left kidney at EI/EE (cm); EI-end inspiration, EE-end expiration. |  |  |  |  |  |

In addition to the global comparison between male and female groups of healthy children, we also performed age and gender-based subgroup analyses, including comparisons between groups MG1 vs. FG1, MG2 vs. FG2, …, and MG5 vs. FG5 as shown in supplementary **Tables S2, S3**,**S4**,**S5 and S6**.

Figure 4 illustrates the measurements of those five features in Table 3 across all age groups for healthy children for male and female groups separately. It is obvious that object parameter measurements for MG3 and FG3 groups behave differently from those of other subgroups. For example, the measurements of M1, M3, and M9 are greater in the female group FG3 compared to those in the male group MG3, which differs from what is observed in the other age-based subgroups where the male groups have higher values. This phenomenon was also observed for other object parameter measurements such as M16, M29, and M41.

**Figure 4.**
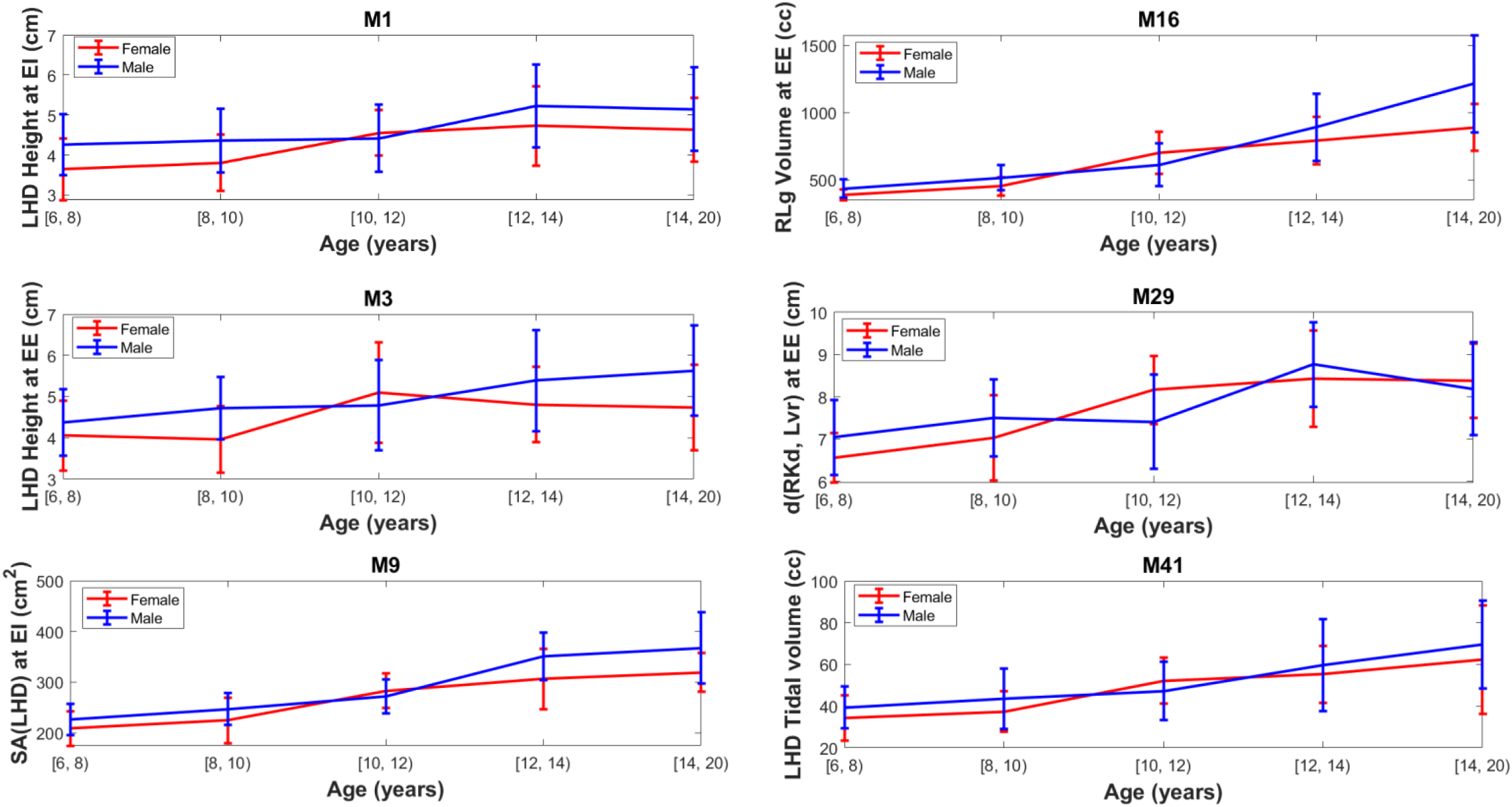
Salient object parameter measurements within age-based subgroups between male and female groups of healthy children. LHD – left hemi-diaphragm, RLg – right lung, RKd – right kidney, Lvr – liver, SA(A) – surface area of object A (in cm^2^), d(A,B) – distance between centroids of object A and B (in cm), EE – end expiration, EI – end inspiration.

BLVee (derived from QdMRI) and FRC (derived from PFT) were highly correlated with a correlation coefficient of 0.86 as shown in Figure 5 in a subset of 51 healthy children from the VGC study cohort.

**Figure 5.**
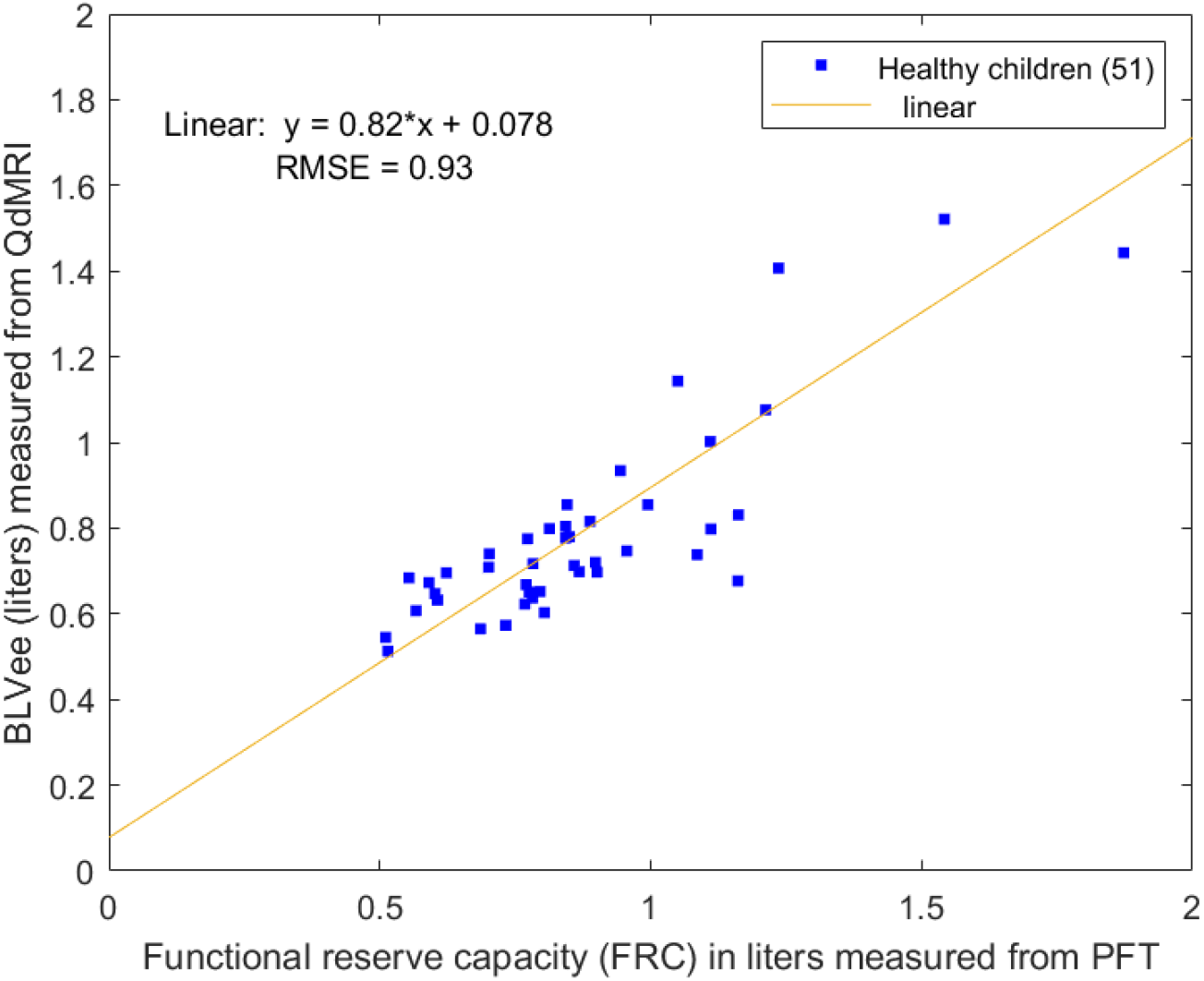
Correlation between QdMRI and pulmonary function testing measurements in a subset of 51 healthy children from the VGC study cohort. PFT – pulmonary function test.

### Application of VGC measurements/ models for assessment of the TIS patient cohort

#### 1) Comparisons of pre-operative vs. post-operative TIS patients

Table 4 illustrates the salient measurements which showed significant differences between pre-operative and post-operative groups of TIS patients, where age-correction was performed on TIS patients before surgery. These measurements generally increased after surgery including left and right hemi-diaphragm heights at EI and EE, left and right lung tidal volumes, and hemi-chest wall tidal volumes, certain architectural parameters as listed, and the motion of specific regions of the hemi-diaphragms such as RHD-CMR (the center and middle region of right hemi-diaphragm). All other parameter measurements (among the 103) did not show significant increases from before to after surgery.

**Table 4.** Salient measurements (mean and SD) showing significant differences between pre-operative and post-operative TIS patients.

| Measurement | M1 | M2 | M3 | M4 | M6 | M55 | M66 | M72 |
| --- | --- | --- | --- | --- | --- | --- | --- | --- |
| **TIS pre-op** | 3.19<br>1.00 | 3.65<br>1.18 | 3.38<br>1.53 | 3.55<br>1.09 | 11.78<br>2.66 | 8.3<br>1.69 | 83.42<br>30.58 | 8.36<br>1.73 |
| **TIS post-op** | 3.73<br>1.19 | 3.96<br>1.29 | 4.11<br>2.04 | 3.96<br>1.15 | 12.42<br>2.70 | 8.64<br>1.73 | 95.28<br>28.75 | 8.73<br>1.72 |
|  | 0.001 | 0.030 | 0.001 | 0.006 | 0.010 | 0.025 | 0.031 | 0.018 |

**Table 4 (continued).**
| Measurement | M79-<br>'RHD<br>-PR' | M81-<br>'RHD<br>-MR' | M82-<br>'RHD<br>-CR' | M87-<br>'RHD<br>-PMR' | M88-<br>'RHD-PCR' | M90-<br>'RHD<br>-CMR' | M37 | M38 | M39 | M40 |
| --- | --- | --- | --- | --- | --- | --- | --- | --- | --- | --- |
| **TIS pre-op** | 4.40<br>1.82 | 3.08<br>2.28 | 4.66<br>1.75 | 3.44<br>2.92 | 4.79<br>2.19 | 3.33<br>2.39 | 43.28<br>22.36 | 65.15<br>37.83 | 27.86<br>15.82 | 32.41<br>20.87 |
| **TIS post-op** | 6.28<br>5.61 | 4.88<br>4.74 | 6.38<br>4.94 | 6.08<br>7.53 | 7.07<br>6.85 | 4.89<br>4.13 | 50.93<br>27.59 | 79.25<br>45.40 | 32.65<br>17.95 | 40.42<br>21.44 |
|  | 0.034 | 0.024 | 0.030 | 0.031 | 0.037 | 0.032 | 0.015 | 0.004 | 0.022 | 0.006 |

M1/M2 - left/right diaphragm height at EI (cm); M3/M4 - left/right diaphragm height at EE(cm); P10 – right lung height at EI(cm); M55/M72 – distance of centroids of (left, right lungs) at EI/EE (in cm);
M66 - angle between the centroids of the right lung and left hemi-diaphragm with the dome of the right hemi-diaphragm as the angle vertex at EE;
M79 - RHD-PR, the average velocity in right hemi-diaphragm posterior region (mm/sec)
M81 - RHD-MR, the average velocity in right hemi-diaphragm medial region (mm/sec)
M82 - RHD-CR, the average velocity in right hemi-diaphragm central region (mm/sec)
M87 - RHD-PMR, the average velocity in right hemi-diaphragm posterior-medial region (mm/sec)
M88 - RHD-PCR, the average velocity in right hemi-diaphragm posterior-central region (mm/sec)
M90 - RHD-CMR, the average velocity in right hemi-diaphragm central-medial region (mm/sec)
M37/M38 - left/right lung tidal volume (in cc);

In Table 5, we specifically listed those volumes and tidal volumes that were significantly different between pre-operative and post-operative TIS patients. In particular, the volumes of the left and right lungs at EI and EE (M13, …, M16) and the tidal volumes of the left and right lungs and left and right hemi-chest walls (M37, …,M40) increased after surgery.

**Table 5.** Volume (in cc) comparisons between pre-operative and post-operative TIS patients with age correction. Mean and SD are reported.

|  | M13 | M14 | M15 | M16 | M37 | M38 | M39 | M40 |
| --- | --- | --- | --- | --- | --- | --- | --- | --- |
| **TIS pre-op** | 322.22<br>168.77 | 432.28<br>244.69 | 278.94<br>154.25 | 367.13<br>213.42 | 43.27<br>22.36 | 65.15<br>37.83 | 27.86<br>15.82 | 32.41<br>20.87 |
| **TIS post-op** | 256.34<br>149.12 | 378.14<br>232.60 | 205.41<br>129.32 | 298.89<br>195.73 | 51.17<br>27.24 | 79.25<br>45.40 | 32.65<br>17.95 | 40.42<br>21.44 |
|  | <0.001 | <0.001 | <0.001 | <0.001 | 0.012 | 0.004 | 0.02 | 0.006 |

M13/M14- left/right lung volume at EI; M15/M16 -left/ right lung volume at EE; M37/M38- left/right lung tidal volume; M39/M40 -Left/right chest wall tidal volume. All volumes with the unit in cc.

#### 2) Comparisons of pre-operative TIS patient deviations from normal vs. post-operative TIS patient deviations from normal

Measurements from the VGC study healthy child cohort provide reference standard measurements for comparing TIS patients before and after surgery.

Table 6a shows Salient measurements (mean and SD using Mahalanobis distance (MD)) showing significant differences between TIS patients (pre-operative or post-operative) and healthy children less than 6 years. For normal children less than 6 years, extrapolation via a linear regression model was performed to estimate the measurements. Table 6b shows the results of comparisons between TIS patients (before or after surgery) vs. healthy children when the healthy children equal or elder than 6 years.

**Table 6a.** Salient measurements (mean and SD using Mahalanobis distance (MD)) showing significant differences between TIS patients (pre-operative or post-operative) and healthy children (<6 years). P values comparing MD (pre-operative TIS vs. healthy) vs. MD (post-operative TIS vs. healthy) using paired t-testing are reported.

|  |  |  |  |  |  |
| --- | --- | --- | --- | --- | --- |
|  | M1 | M3 | M13 | P15 | M16 |
| **MD (pre-operative TIS vs. healthy)** | -0.87<br>2.91 | -1.36<br>2.33 | 2.09<br>2.23 | 2.56<br>2.48 | 1.87<br>2.56 |
| **MD (post-operative TIS vs. healthy)** | 0.53<br>3.90 | 0.44<br>4.69 | 1.35<br>2.79 | 1.66<br>3.03 | 1.16<br>3.15 |
| **P value** | 0.01 | 0.02 | 0.01 | 0.00 | 0.02 |
| M1/M3- left hemi-diaphragm height at EI/EE; M13/M15- LLVei/LLVee; M16-RLVee. |  |  |  |  |  |

**Table 6b.** Salient measurements (mean and SD using Mahalanobis distance (MD)) showing significant differences between TIS patients (pre-operative or post-operative) and healthy children (≥6 years). P values comparing MD (pre-operative TIS vs. healthy) vs. MD (post-operative TIS vs. healthy) using paired t-testing are reported.

|  |  |  |  |  |
| --- | --- | --- | --- | --- |
|  | M1 | M15 | M55 | M72 |
| **MD (pre-operative TIS vs. healthy)** | -0.82<br>2.88 | 1.85<br>3.15 | -1.83<br>1.97 | -2.11<br>2.45 |
| **MD (post-operative TIS vs. healthy)** | -0.10<br>3.14 | 1.51<br>2.97 | -1.02<br>2.71 | -0.90<br>3.57 |
| **P value** | 0.04 | 0.03 | 0.00 | 0.00 |
| M1- left hemi-diaphragm height at EI; M15-LLVee; M55/M72 distance of lungs (left, right) at EI/EE. |  |  |  |  |

We observed that the MD (i.e., deviation from normality) for left hemi-diaphragm height at EI and EE, left and right lung volumes at EE, and left lung volume at EI in younger TIS patients decreased after treatment as shown in Table 6a. Also, measurements of left diaphragm height at EI (M1); LLVee(M15), distance of lungs (left, right) at EI/EE (M55/M72) became closer to normal in older TIS patients after treatment as shown in Table 6b.

## 3. Discussion

Quantitative measurements obtained from dynamic MRI in the VGC study of healthy children have provided new insights into the normal maturation process of healthy children in terms of thoracoabdominal structure and function, and can serve as useful reference standards for quantitative assessment of abnormalities in regional respiratory function in pediatric or adults patients with disorders such as TIS.

### Maturation of normal children

Some interesting findings were observed in the comparisons of the male and female groups of healthy children globally and among specific age subgroups. For example, in global comparisons (94 male vs. 97 female), diaphragm height at EI or EE, diaphragm surface area, and the distance between kidney centroids (at EI and EE, respectively) were larger in males than in females, as shown in **Table 3**.

In age-based sub-group comparisons, such as between males and females of ages 6-8 years, left hemi-diaphragm height at EI and EE were also larger in males compared to females as shown in **Table S2**. Interestingly, we observed that comparisons of parameter measurements between male and female children of ages 10-12 years seemed to behave differently from male vs. female in the other age-based sub-group comparisons. For example, left hemi-diaphragm height at EI is lower for male group (10-12 years) than that for female group (10-12 years), which is inversed in other sub-group (6-8 year group, 8-10 year group, 12-14 year group, group with age above 14 years) comparison. Similarly, the distance between right kidney centroid and the liver centroid, Right hemi-diaphragm tidal volume of male group (10-12 years) with smaller values than female group (10-12 years). 10-12 years for females is a special growth range, the growth rate for females continues to be greater than males between 10 and 12 years of age. Previous research [44, 45] also show the similar observation that after 13 years of age, the growth spurt of females generally is completed and the growth spurt of males is in its early phase. Those publications [44, 45] can only give a global evaluation of growth, however, QdMRI first time gives deeper insight via regional measurements between male and female groups.

The comparison of male vs. female in sub-group 5 (with the age above 14 years) in **Table S6** shows more features than any other sub-groups to distinguish male and female groups which might mean the different between adolescent male and female will be more significant than the younger children. Those features including lung and diaphragm height measurement, (M3-M8), and diaphragm surface area (M9-M12), lung volumes (M13-M16), distance of right and left kidney at EI, and also architectures of distance between left and right lungs, distance between lung centroid and diaphragm dome, and lung volumes at EI and EE as shown in **Table S6**.

### PFT and QdMRI parameters

Within the subset of 51 healthy children who underwent QdMRI and pulmonary function testing (PFT), we observed a high correlation of 0.86 between bilateral lung volume at end-expiration (BLVee) derived from QdMRI and the functional residual capacity (FRC) derived from PFT. BLVee does not equal to FRC. Many factors might contribute the inequality between them. There might be much more natural variation in the QdMRI measures than PFT which could also lead to difference between them. Some other factors include the difficulties in checking blood effects on image-based volume estimation and the accurate separation of the lung and upper airway.

QdMRI measurements might be affected by the segmentation challenge on MRI from the image quality, motion and blood flow effect. Yet, QdMRI provides additional value compared to PFT alone, as the latter cannot be used in patients who are very young or unable to cooperate and does not provide regional respiratory functional information, including for example the contribution of each lung, hemi-chest wall, or hemi-diaphragm to respiratory function. In addition, QdMRI allows one to analyze the properties of multiple objects of interest beyond the lungs alone including those related to architecture and dynamics simultaneously.

### Quantitative assessment of TIS patients using the VGC system

TIS patients are usually treated via surgery, such as VEPTR, to provide much more space for lung to extend and grow. It is crucial to compare the patient before and after surgery in order to ensure and adjust continue treatment plan. Besides the Cobb angle and PFT and other questionnaire limited approaches check on patients after surgery, there are no more practical and quantitative methods to evaluate the treatment effects and provide useful information for treatment. Some young TIS patients it is almost impossible to ask them to collaborate any on PFT. QdMRI based approach allows the patients free breathing image acquisition and can be a practical approach to quantitatively evaluate the specific organs motion, architure, and VGC system providing the normal reference data which can furthermore improve the understanding of those parameters by checking the closeness of them before surgery to normal, and after surgery to normal by using mahalanobis distance (MD). Such a large normal children measurement database is a unique and allows multiple analysis purposes. We believe given any measurement/factor, the normal value for it should be in a “normal” range instead of a single value, in other words, the normal measurement means a group of normal subjects who provide the useful information as a reference.

TIS patients usually suffer from the deform of spine which limit the normal grow or mutation. When we compare TIS patient before and after surgery using age-correction as shown in **Table 4**, where all measurements are significantly increased, including diaphragm height, left and right lung distance, and the diaphragm motion, and more importantly, the tidal lung volume and chest wall tidal volume are significantly improved, which means the thoracic dynamics and function improved after surgery. Those results are valuable in practice. There is a concern in pediatric surgery practice that rib-based fixation may limit chest wall motion in TIS. Several studies have been published to assess that concern [46-48]. But generally, those publications did not solid answer the concern and those techniques can be barely used on our TIS patients. Those publications did not directly compare the same patients before and after surgery; and also, very limited normal subjects are used, such as only 12 or 9 normal individuals in [46-48] or even only one patient is used [47]. We used quantitative measurements from dynamic MRI using 191 normal children and 49 paired TIS patients which are far beyond the data in any existing literature. We observed results that chest wall tidal volume increases after surgery which suggests that chest wall motion is not impaired in pediatric patients with TIS after rib-based surgery.

We specifically compare the lung volume (M13-M16) and tidal volume (M37-M40) with age correction in Table 5. The lung tidal volume shows significant increase although the lung volume did not show the significant improvement. In other words, although the lung size does not increase significantly (to the corresponding normal children level after surgery), but the lung dynamics and function from tidal volume are significantly improved.

Mahalanobis distance (MD) can be used to evaluate the similarity of one TIS subject to the corresponding healthy children subset. The comparison of two MDs with MD1 between TIS pre-op subjects vs. their corresponding normal children comparing and MD2 between TIS post-op subject vs. their corresponding normal children do not behave same for younger TIS patients vs. the older TIS patients.

For the younger TIS patients and their corresponding healthy children (<6 years), left hemi-diaphragm and left lung volumes closer to the normal after surgery as shown in Table 6a. But the 2D height measurements and 3D volume show different properties and left and right sides measurements seem not to be symmetrical. The features of left hemi-diaphragm height at both EI and EE significantly close to the normal, Interestingly, the left lungs at EI and EE also show the same properties. Left lung volume at both EI and EE closer to the normal after surgery, and right lung only at EE (RLVee) closer to the normal. And right hemi-diaphragm height at EI (M2) did not show significant difference, and the right hemi-diaphragm height at EE (M4 with larger MD) is still a little farther from to normal which indicate the right hemi-diaphragm and its motion might still be limited by the TIS after surgery and need to further improve which limited the right lung expending at EI.

For the older TIS patient and the corresponding healthy children (≥ 6 years), new feature of P46, P93, the distance between left and right lung centers are significantly improved and closers to the normal after surgery. The M1 and M14 (the left hemi-diaphragm height at EI and left lung volume at EE) which also shows up in the younger TIS group also show less MDs and are closer to the normal after surgery as shown in **Table 6b**. The right components of right hemi-diaphragm did not show more closeness to the normal after surgery which might indicate that the right hemi-diaphragm height should be further improved which also indicates the left and the right side is not symmetric and the surgery plan might need to adjust more and to allow more on the right hemi-diaphragm motion.

There are some limitations of this study. Currently, the VGC study of healthy children only includes those who are older than 6 years of age, and so used extrapolation instead of real healthy children data sets to estimate normal parameter measurements for comparison with very young TIS patients less than 6 years of age before and after surgery. Another limitation is that data sets were all obtained from one center for both healthy children and TIS patients. Future research may be performed to obtain more QdMRI data from other centers, as well as from patients with disorders other than TIS.

## 4. Conclusions

In this paper, we introduce the virtual growing child (VGC), which includes a large unique database of free-breathing thoracoabdominal 4D dynamic MRI images, object segmentations, and associated quantitative normative reference standard measurements and models obtained from healthy children. This system can be useful to quantify regional respiratory anomalies and facilitate treatment planning and response assessment, as we have demonstrated in the setting of pediatric thoracic insufficiency syndrome (TIS). The VGC database can be useful to advance future AI-based research on dMRI-based object segmentation and analysis.

## Supplementary Tables

Table S1: Quantitative parameter measurement definitions.

Table S2. Salient measurement (mean and SD) comparisons between MG1(22) vs. FG1(20) normal children.

Table S3: Salient measurement (mean and SD) comparisons between male (MG2) vs. female (FG2) 8-10 year old healthy children.

Table S4: Salient measurement (mean and SD) comparisons between male (MG3) vs. female (FG3) 10-12 year old healthy children.

Table S5: Salient measurement (mean and SD) comparisons between male (MG4) vs. female (FG4) 12-14 year old healthy children.

Table S6: Salient measurement (mean and SD) comparisons between male (MG5) vs. female (FG5) 14-18 year old healthy children.

