## Supplemental Table S1 for "Virtual Growing Child (VGC): A general normative comparative system via quantitative dynamic MRI for quantifying pediatric regional respiratory anomalies with application in thoracic insufficiency syndrome (TIS)"

| Table S1. Quantitative measurement definition. | |
|---|---|
| **Measurements** | Notes |
| **M1** | Left diaphragm height at EI (cm) |
| **M2** | Right diaphragm height at EI (cm) |
| **M3** | Left diaphragm height at EE (cm) |
| **M4** | Right diaphragm height at EE (cm) |
| **M5** | Left lung height at EI (cm) |
| **M6** | Right lung height at EI (cm) |
| **M7** | Left lung height at EE (cm) |
| **M8** | Right lung height at EE (cm) |
| **M9** | Left diaphragm surface area at EI ($cm^2$) |
| **M10** | Right diaphragm surface area at EI ($cm^2$) |
| **M11** | Left diaphragm surface area at EE ($cm^2$) |
| **M12** | Right diaphragm surface area at EE ($cm^2$) |
| **M13** | LLVei, left lung volume at end inspiration (cc) |
| **M14** | RLVei, right lung volume at end inspiration (cc) |
| **M15** | LLVee, left lung volume at end expiration (cc) |
| **M16** | RLVee, right lung volume at end expiration (cc) |
| **M17** | distance between RLD dome and liver centroid at EI (cm) |
| **M18** | distance between RLD dome and right kidney centroid at EI (cm) |
| **M19** | distance between Right kidney centroid and liver centroid at EI (cm) |
| **M20** | angle of (M17 and M18) at EI in degree, the angle between the centroids of liver and right kidney with the dome of the right hemi-diaphragm as the angle vertex at EI |
| **M21** | angle of (M17 and M19) at EI in degree, the angle between right hemi-diaphragm dome, and the centroid of right kidney with the centroid of the liver as the angle vertex at EI |
| **M22** | distance between LLD dome and Left kidney centroid at EI (cm) |
| **M23** | distance between LLD dome and Rkidney centroid at EI (cm) |
| **M24** | distance between centroids of Rkidney and Lkidney at EI (cm) |
| **M25** | angle (M22, M23) at EI in degree |
| **M26** | angle (M22, M24) at EI in degree |
| **M27** | distance between RLD dome and liver centroid at EE (cm) |
| **M28** | distance between the dome of right hemi-diaphragm and the centroid of right kidney at EE (cm) |
| **M29** | distance between the centroids of right kidney and liver at EE (cm) |
| **M30** | angle of (M27 and M28) at EE in degree |
| **M31** | angle of (M27 and M29) at EE in degree |
| **M32** | distance between LLD dome to Lkidney centroid at EE (cm) |
| **M33** | distance between LLD dome to Rkidney centroid at EE (cm) |
| **M34** | distance between centroids of Rkidney and Lkidney at EE (cm) |
| **M35** | angle (M32, M33) at EE in degree |
| **M36** | angle (M32, M34) at EE in degree |
| **M37** | LLtv, Left lung tidal volume (cc) |
| **M38** | RLtv, Right lung tidal volume (cc) |
| **M39** | LCWtv, left chest wall tidal volume (cc) |
| **M40** | RCWtv, right chest wall tidal volume (cc) |

| | |
|---|---|
| **M41** | Ldtv, left diaphragm tidal volume (cc) |
| **M42** | Rdtv, right diaphragm tidal volume (cc) |
| **M43** | distance between (RLg, RLD) at EI (cm) |
| **M44** | distance between (RLD, LLD) at EI (cm) |
| **M45** | distance between (RLg, LLD) at EI (cm) |
| **M46** | angle (M43, M44), the angle between the centroids of the right lung and left hemi-diaphragm with the dome of the right hemi-diaphragm as the angle vertex at EI |
| **M47** | angle (M43, M45), the angle between the domes of left and right hemi-diaphragams the centroid of right lung as the angle vertex at EI |
| **M48** | distance between (LLg, RLD) at EI (cm) |
| **M49** | distance between (RLD, LLD) at EI (cm) |
| **M50** | distance between (LLg, LLD) at EI (cm) |
| **M51** | angle (M48, M49), the angle between the centroids of the left lung and left hemi-diaphragm with the dome of the right hemi-diaphragm as the angle vertex at EI |
| **M52** | angle (M48, M50), the angle between the centroids of the right and left hemi-diaphragms with the center of left lung as the angle vertex at EI |
| **M53** | distance between (RLg, LLD) at EI (cm) |
| **M54** | distance between (LLg, LLD) at EI (cm) |
| **M55** | distance between (LLg, RLg) at EI (cm) |
| **M56** | angle (M53, M54), the angle between the centroids of the left and right lungs with the dome of the left hemi-diaphragm as the angle vertex at EI |
| **M57** | angle (M53, M55), the angle between the centroids of the left lung and left hemi-diaphragms with the center of right lung as the angle vertex at EI |
| **M58** | distance between (RLg, RLD) EI (cm) |
| **M59** | distance between (RLD, LLg) at EI (cm) |
| **M60** | distance between (LLg, RLg) at EI (cm) |
| **M61** | angle (M58, M59), the angle between the centroids of the left and right lungs with the dome of the right hemi-diaphragm as the angle vertex at EI |
| **M62** | angle (M58, M60), the angle between the centroids of the left lung and right hemi-diaphragms with the center of right lung as the angle vertex at EI |
| **M63** | distance between (RLg, RLD) at EE (cm) |
| **M64** | distance between (RLD, LLD) at EE (cm) |
| **M65** | distance between (LLD, RLg) at EE (cm) |
| **M66** | Distance between centroid of right lung and the left hemi-diaphragm dome at EE (cm) |
| **M67** | angle (M63, M65), the angle between the centroids of the right and left hemi-diaphragms with the center of right lung as the angle vertex at EE |
| **M68** | distance between (LLg, RLD) at EE (cm) |
| **M69** | the angle between the centroids of the left lung and left hemi-diaphragm with the dome of the right hemi-diaphragm as the angle vertex at EE |
| **M70** | the angle between the centroids of the right and left hemi-diaphragms with the center of left lung as the angle vertex at EE |
| **M71** | distance between (LLD, LLg) at EE (cm) |
| **M72** | distance between (LLg, RLg) at EE (cm) |
| **M73** | The angle between the centroid of right lung and the dome of the right hemi-diaphragm with left hemi-diaphragm dome as the angle vertex at EE |
| **M74** | the angle between the centroids of the left lung and left hemi-diaphragms with the center of right lung as the angle vertex at EE |
| **M75** | distance between (LLg, RLg) at EE (cm) |
| **M76** | the angle between the centroids of the left and right lungs with the dome of the right hemi-diaphragm as the angle vertex at EE |

| | |
|---|---|
| **M77** | the angle between the centroids of the left lung and right hemi-diaphragms with the center of right lung as the angle vertex at EE |
| **M78** | RHD-AR, the average velocity in right hemi-diaphragm anterior region (mm/sec) |
| **M79** | RHD-PR, the average velocity in right hemi-diaphragm posterior region (mm/sec) |
| **M80** | RHD-LR, the average velocity in right hemi-diaphragm later region (mm/sec) |
| **M81** | RHD-MR, the average velocity in right hemi-diaphragm medial region (mm/sec) |
| **M82** | RHD-CR, the average velocity in right hemi-diaphragm central region (mm/sec) |
| **M83** | RHD-ALR, the average velocity in right hemi-diaphragm anterior-lateral region (mm/sec) |
| **M84** | RHD-AMR, the average velocity in right hemi-diaphragm anterior-medial region (mm/sec) |
| **M85** | RHD-ACR, the average velocity in right hemi-diaphragm anterior-medial region (mm/sec) |
| **M86** | RHD-PLR, the average velocity in right hemi-diaphragm posterior-lateral region (mm/sec) |
| **M87** | RHD-PMR, the average velocity in right hemi-diaphragm posterior-medial region (mm/sec) |
| **M88** | RHD-PCR, the average velocity in right hemi-diaphragm posterior-central region (mm/sec) |
| **M89** | RHD-CLR, the average velocity in right hemi-diaphragm central-lateral region (mm/sec) |
| **M90** | RHD-CMR, the average velocity in right hemi-diaphragm central-medial region (mm/sec) |
| **M91** | LHD-AR, the average velocity in left hemi-diaphragm anterior region (mm/sec) |
| **M92** | LHD-PR, the average velocity in left hemi-diaphragm posterior region (mm/sec) |
| **M93** | LHD-LR, the average velocity in left hemi-diaphragm later region (mm/sec) |
| **M94** | LHD-MR, the average velocity in left hemi-diaphragm medial region (mm/sec) |
| **M95** | LHD-CR, the average velocity in left hemi-diaphragm central region (mm/sec) |
| **M96** | LHD-ALR, the average velocity in left hemi-diaphragm anterior-lateral region (mm/sec) |
| **M97** | LHD-AMR, the average velocity in left hemi-diaphragm anterior-medial region (mm/sec) |
| **M98** | LHD-ACR, the average velocity in left hemi-diaphragm anterior-medial region (mm/sec) |
| **M99** | LHD-PLR, the average velocity in left hemi-diaphragm posterior-lateral region (mm/sec) |
| **M100** | LHD-PMR, the average velocity in left hemi-diaphragm posterior-medial region (mm/sec) |
| **M101** | LHD-PCR, the average velocity in left hemi-diaphragm posterior-central region (mm/sec) |
| **M102** | LHD-CLR, the average velocity in left hemi-diaphragm central-lateral region (mm/sec) |
| **M103** | LHD-CMR, the average velocity in left hemi-diaphragm central-medial region (mm/sec) |
| **EI, end inspiration; EE, end expiration; RLg, right lung; LLg, left lung; RLD, right lateral hemi-diaphragm; LLD, left lateral hemi-diaphragm.** | |
