## Supplemental Table S2 for "Virtual Growing Child (VGC): A general normative comparative system via quantitative dynamic MRI for quantifying pediatric regional respiratory anomalies with application in thoracic insufficiency syndrome (TIS)"

**Table S2. Salient measurement (mean and SD) comparisons between MG1(22) vs. FG1(20) normal children.**

| | M1 | M5 | M7 | M9 | M13 | M14 | M16 | M29 | M40 |
|---|---|---|---|---|---|---|---|---|---|
| **Male** | 4.23<br>0.75 | 15.18<br>0.90 | 14.06<br>0.94 | 227.33<br>30.94 | 382.65<br>55.96 | 527.75<br>78.97 | 432.16<br>66.67 | 7.10<br>0.91 | 46.77<br>12.84 |
| **Female** | 3.60<br>0.77 | 14.50<br>1.00 | 13.38<br>0.94 | 207.03<br>33.29 | 348.31<br>47.06 | 467.29<br>49.87 | 384.86<br>40.14 | 6.54<br>0.57 | 35.76<br>9.56 |
| **P value** | 0.01 | 0.03 | 0.02 | 0.05 | 0.04 | 0.01 | 0.01 | 0.02 | <0.001 |

M1 - Left hemi-diaphragm at EI (cm)
M5 - Left lung height at EI (cm)
M7 - Left lung height at EE (cm)
M9 - Left hemi-diaphragm surface area at EI ($cm^2$)
M13 - LLVei, left lung volume at EI (cc)
M14 - RLVei (cc), right lung volume at EI (cc)
M16 - RLVee (cc), right lung volume at EE (cc)
M29 - Distance between centroids of right kidney and liver at EE (cm)
M40 - RCWtv, right chest wall tidal volume (cc)
EI - End inspiration
EE - End expiration
Note:
p value (M5 vs. M7) < 0.001 for MG1 or FG1
p value (M14 vs. M16) <0.001 for MG1 or FG1
p value (M13 vs. M14) < 0.001 for MG1 or FG1
