## Supplemental Table S3 for "Virtual Growing Child (VGC): A general normative comparative system via quantitative dynamic MRI for quantifying pediatric regional respiratory anomalies with application in thoracic insufficiency syndrome (TIS)"

**Table S3. Salient measurement (mean and SD) comparisons between MG2(20) vs. FG2(16) normal children.**

| | M1 | M3 | M7 | M16 | M34 | M66 | M73 |
|---|---|---|---|---|---|---|---|
| **Male** | 4.39<br>0.81 | 4.74<br>0.78 | 15.31<br>1.22 | 522.51<br>89.18 | 9.66<br>0.68 | 10.64<br>1.82 | 69.65<br>10.90 |
| **Female** | 3.86<br>0.69 | 4.01<br>0.82 | 14.49<br>1.10 | 458.73<br>63.25 | 9.03<br>0.77 | 9.35<br>1.97 | 80.85<br>13.31 |
| **P value** | 0.043 | 0.009 | 0.043 | 0.021 | 0.013 | 0.049 | 0.009 |

M1 - Left hemi-diaphragm height at EI (cm)
M3 - Left hemi-diaphragm height at EE (cm)
M7 - Left lung height at EE (cm)
M16 - RLVee, right lung volume at EE (cc)
M34 - Distance between centroids of Rkidney and Lkidney at EE (cm)
M66 – Distance between centroid of right lung and the left hemi-diaphragm dome at EE (cm)
M73 - The angle between the centroid of right lung and the dome of the right hemi-diaphragm with left hemi-diaphragm dome as the angle vertex at EE
EI - End inspiration
EE - End expiration
Note:
P value (M1 vs. M3) =0.02 for MG2
P value (M1 vs. M3) = 0.33 for FG2
