## Supplemental Table S4 for "Virtual Growing Child (VGC): A general normative comparative system via quantitative dynamic MRI for quantifying pediatric regional respiratory anomalies with application in thoracic insufficiency syndrome (TIS)"

**Table S4. Salient measurement (mean and SD) comparisons between MG3(12) vs. FG3(17) normal children.**

| | M5 | M6 | M7 | M20 | M21 | M28 | M29 | M38 | M42 | M46 | M47 | M61 | M62 | M71 |
|---|---|---|---|---|---|---|---|---|---|---|---|---|---|---|
| **Male** | 16.91<br>1.96 | 16.78<br>1.47 | 15.72<br>1.87 | 34.74<br>10.88 | 113.16<br>21.63 | 11.38<br>1.99 | 7.41<br>1.11 | 120.80<br>33.38 | 77.38<br>20.23 | 86.85<br>28.45 | 68.19<br>30.35 | 59.39<br>13.62 | 96.87<br>17.57 | 5.83<br>1.58 |
| **Female** | 18.87<br>1.69 | 18.13<br>1.71 | 17.57<br>1.81 | 26.75<br>8.84 | 132.84<br>15.78 | 12.77<br>1.13 | 8.16<br>0.80 | 148.49<br>35.32 | 96.94<br>24.52 | 109.95<br>23.07 | 41.67<br>22.69 | 73.88<br>14.08 | 78.41<br>17.98 | 6.99<br>1.34 |
| **P value** | 0.01 | 0.04 | 0.01 | 0.04 | 0.01 | 0.02 | 0.04 | 0.04 | 0.03 | 0.02 | 0.01 | 0.01 | 0.01 | 0.04 |

M5 – Left lung height at EI (cm)
M6- Right lung height at EI (cm)
M7- Left lung height at EE (cm)
M20 - the angle between the centroids of liver and right kidney with the dome of the right hemi-diaphragm as the angle vertex at EI
M21 - the angle between right hemi-diaphragm dome, and the centroid of right kidney with the centroid of the liver as the angle vertex at EI
M28 – distance between the dome of right hemi-diaphragm and the centroid of right kidney at EE (cm)
M29 - distance between the centroids of right kidney and liver at EE (cm)
M38 - right lung tidal volume (cc)
M42 – RDtv, right hemi-diaphragm tidal volume (cc)
M46 - angle (M43, M44), the angle between the centroids of the right lung and left hemi-diaphragm with the dome of the right hemi-diaphragm as the angle vertex at EI
M47 - angle (M43, M45), the angle between the domes of left and right hemi-diaphragms with the centroid of right lung as the angle vertex at EI
M61 – angle (M58, M59), the angle between the centroids of left and right lungs with the dome of right hemi-diaphragm as the angle vertex at EI
M62 – the angle between the centroid of left lung and the right hemi-diaphragm dome with the centroid of right lung as the angle vertex at EI
M71 – distance between the dome of left hemi-diaphragm and the centroid of left lung (LLD, LLg) at EE(cm)
