## Supplemental Table S5 for "Virtual Growing Child (VGC): A general normative comparative system via quantitative dynamic MRI for quantifying pediatric regional respiratory anomalies with application in thoracic insufficiency syndrome (TIS)"

**Table S5. Salient measurement (mean and SD) comparisons between MG4(18) vs. FG4(14) normal children.**

| | M9 | M10 | M11 | M24 | M34 | M40 | M76 |
|---|---|---|---|---|---|---|---|
| **Male** | 350.57<br>47.14 | 372.74<br>65.89 | 330.34<br>56.72 | 11.53<br>0.86 | 11.31<br>0.88 | 78.13<br>35.01 | 74.04<br>16.75 |
| **Female** | 305.49<br>57.02 | 325.68<br>59.53 | 282.91<br>64.21 | 10.45<br>0.95 | 10.34<br>0.84 | 53.32<br>28.16 | 62.90<br>12.93 |
| **P value** | 0.020 | 0.045 | 0.034 | 0.002 | 0.004 | 0.039 | 0.049 |

**M9 - left hemi-diaphragm surface area at EI in $cm^2$**
**M10 - right hemi-diaphragm surface area at EI in $cm^2$**
**M11 - left hemi-diaphragm surface area at EE in $cm^2$**
**M24 -** distance **between centroids of Rkidney and Lkidney at EI in cm**
**M34 - distance between centroids of Rkidney to Lkidney at EE in cm**
**M40 - RCWtv in cc**
**M76, angle (RLg, RLD), and (RLD, LLg) at EE**
