## Supplemental Table S6 for "Virtual Growing Child (VGC): A general normative comparative system via quantitative dynamic MRI for quantifying pediatric regional respiratory anomalies with application in thoracic insufficiency syndrome (TIS)"

**Table S6. Salient measurement (mean and SD) comparisons between MG5 (22) vs. FG5(30) normal children.**

| | M2 | M5 | M6 | M7 | M8 | M9 | M10 | M11 | M12 | M13 | M14 | M15 | M16 |
|---|---|---|---|---|---|---|---|---|---|---|---|---|---|
| **Male** | 5.63<br>1.09 | 21.47<br>2.55 | 20.99<br>2.43 | 20.61<br>2.70 | 20.07<br>2.58 | 367.34<br>70.40 | 397.95<br>65.81 | 342.45<br>67.66 | 357.13<br>60.63 | 1084.22<br>312.63 | 1347.00<br>382.45 | 995.22<br>308.90 | 1215.69<br>362.63 |
| **Female** | 47.97<br>9.94 | 19.89<br>1.91 | 19.44<br>1.85 | 18.62<br>1.61 | 17.74<br>1.53 | 319.89<br>38.29 | 352.78<br>50.20 | 297.51<br>43.24 | 319.31<br>44.07 | 808.35<br>187.41 | 1029.76<br>208.56 | 705.83<br>158.95 | 892.78<br>177.87 |
| **P value** | 0.006 | 0.013 | 0.012 | 0.002 | 0.000 | 0.003 | 0.007 | 0.005 | 0.012 | 0.000 | 0.000 | 0.000 | 0.000 |

**M2 - left hemi-diaphragm height at EE in cm**
**M5 - left lung height at EI in cm**
**M6 - right lung height at EI in cm**
**M7 - left lung height at EE in cm**
**M8 - right lung height at EE in cm**
**M9 - left hemi-diaphragm surface area at EI in $cm^2$**
**M10 - right hemi-diaphragm surface area at EI in $cm^2$**
**M11 - left hemi-diaphragm surface area at EE in $cm^2$**
**M12 - right hemi-diaphragm surface area at EE in $cm^2$**
**M13- LLVei in cc**
**M14 - RLVei in cc**
**M15 - LLVee in cc**
**M16 - RLVee in cc**

**Table S6. (continue).**

| | M24 | M25 | P30 | M31 | M34 | M55 | M65 | M68 | M72 |
|---|---|---|---|---|---|---|---|---|---|
| **Male** | 12.05<br>0.99 | 48.40<br>7.49 | 33.38<br>6.55 | 115.00<br>15.45 | 11.94<br>1.05 | 13.06<br>1.39 | 13.91<br>2.35 | 14.70<br>2.43 | 13.07<br>1.27 |
| **Female** | 10.97<br>0.83 | 43.34<br>5.85 | 28.85<br>7.13 | 127.16<br>13.75 | 10.76<br>0.74 | 12.15<br>1.08 | 12.46<br>2.28 | 13.08<br>1.91 | 12.08<br>1.03 |
| **P value** | 0.000 | 0.009 | 0.023 | 0.004 | 0.000 | 0.011 | 0.030 | 0.010 | 0.003 |

**M24 -side length of Rkidney to Lkidney at EI (cm)**
**M25- angle (M22, M23) at EI in degree**
**M30- angle of (M27 and M28) at EE in degree**
**M31-angle of (M27 and M29) at EE in degree**
**M38 – distance between centroids of Rkidney and Lkidney at EE(cm)**
**M55 distance between (LLg, RLg) at EI (cm)**
**M65, distance between (LLD, RLg) at EE (cm)**
**M68, distance between (LLg, RLD) at EE (cm)**
**M72, distance between (LLg, RLg) at EE (cm)**
